# A discrete-time survival model for porcine epidemic diarrhea virus

**DOI:** 10.1101/2022.06.03.494708

**Authors:** Parker Trostle, Cesar A. Corzo, Brian J. Reich, Gustavo Machado

## Abstract

Since the arrival of porcine epidemic diarrhea virus (PEDV) in the United States in 2013, elimination and control programs have had partial success. The dynamics of its spread are hard to quantify, though previous work has shown that local transmission and the transfer of pigs within production systems are most associated with the spread of PEDV.

Our work relies on the history of PEDV infections in a region of the southeastern United States. This infection data is complemented by farm-level features and extensive industry data on the movement of both pigs and vehicles. We implement a discrete-time survival model and evaluate different approaches to modeling the local-transmission and network effects.

We find strong evidence in that the local-transmission and pig-movement effects are associated with the spread of PEDV, even while controlling for seasonality, farm-level features, and the possible spread of disease by vehicles. Our fully Bayesian model permits full uncertainty quantification of these effects. Our farm-level out-of-sample predictions have a receiver-operating characteristic area under the curve (AUC) of 0.779 and a precision-recall AUC of 0.097. The quantification of these effects in a comprehensive model allows stakeholders to make more informed decisions about disease prevention efforts.

## 1 Introduction

Porcine epidemic diarrhea virus (PEDV) is an economically relevant endemic disease for the U.S. swine industry (Lee 2015). The “puzzle” behind the control and dissemination of PEDV involves complex disease epidemiology from which we still have limited knowledge about its dissemination dynamics and the impact of available control measures (Galvis, Jones, et al. 2022; VanderWaal et al. 2018). Therefore, little has been discovered regarding how PEDV spreads spatially and temporally, which may be directly related to the failure of regional efforts focused on the control and eradication of these diseases (Alvarez et al. 2016; Machado et al. 2019; Niederwerder and Hesse 2018). Tracking the spread of PEDV and porcine reproductive and respiratory syndrome virus (PRRSV) through complex and integrated production systems remains extremely challenging (Machado et al. 2019; VanderWaal et al. 2018).

Swine disease surveillance is used to capture and describe new pathogen circulation, but it generally has limited capabilities to inform about the geographic sources of new infections (Galvis, Jones, et al. 2022; Machado et al. 2019). In addition, most models for disease spread focus either on intrinsic or extrinsic forces driving spread dynamics, which may compromise our ability to fully capture hidden aspects influencing these viruses traveling throughout a complex agricultural landscape and, consequently, our ability to respond in a timely manner. PEDV after its rapid spread to 50% of the U.S. commercial sow farms in 2013 continues to circulate today in U.S. farms (Beam et al. 2015). Since PEDV started to circulate in North America, the swine industry has initiated regional disease control and elimination efforts (Perri et al. 2019).

Despite limited knowledge about the geographic distribution of PEDV, several associated routes of between-farm spread have been identified (Jang et al. 2021; Machado et al. 2019; VanderWaal et al. 2018). With the current knowledge and data available, the movement of animals has been most strongly associated with the propagation of PEDV, particularly the movement of infected replacement gilts (Machado et al. 2019; VanderWaal et al. 2018). In addition, windborne or local spread, such as virus spillover between nearby farms (Alonso et al. 2014; Alvarez et al. 2016; Beam et al. 2015), and contaminated fomites (Lowe et al. 2014) have been demonstrated to be further sources of PEDV propagation. Previous work has shown that the dominant transmission of PEDV appears to be similar to PRRSV, with local spatial spread among neighboring farms and animal movement accounts for 75% of transmission events (Galvis, Corzo, Prada, et al. 2022b; Galvis, Jones, et al. 2022; Machado et al. 2019; VanderWaal et al. 2018).

There have been many proposed modeling approaches in the literature to quantify the dynamics driving the transmission of PEDV or PRRSV. Perhaps the most popular approach is the use of compartmental models and simulations based on those models (Hasahya et al. 2021; Jeong et al. 2014; Murai et al. 2018; Phoongurn et al. 2019; Thakur, Revie, et al. 2015; Thakur, Sanchez, et al. 2015; VanderWaal et al. 2018). Some use Approximate Bayesian Computing combined with compartmental models for parameter estimation and prediction (Galvis, Corzo, and Machado 2022; Galvis, Corzo, Prada, et al. 2022b; Galvis, Jones, et al. 2022; Jones et al. 2021). The advantage of this approach is that the model for the observed data is the same as the proposed theoretical model. However, fitting compartmental models necessitates running many simulations, and this requires a lot of time even for simple models. This may be why many other methods have been considered, such as generalized linear models (Arruda et al. 2017; Haredasht et al. 2017; Lee et al. 2017), spatial clustering (Alvarez et al. 2016; Tousignant et al. 2015), survival analysis (Furutani et al. 2019; Holtkamp et al. 2010), Bayesian phylogenic models (Jara et al. 2021; Makau et al. 2021), and machine-learning approaches (Machado et al. 2019; Silva et al. 2019; Sykes et al. 2021).

We propose using a generalized mixed-effects model as in Haredasht et al. 2017 and Arruda et al. 2017, though we interpret our findings in a survival-analysis context as in Holtkamp et al. 2010 and Furutani et al. 2019. We have access to a comprehensive dataset on animal movements, vehicle movements, and farm-level features, and we are able to fit our model to both productions as well as growth farms. There are many advantages to our approach. The first advantage is that our model is able to incorporate multiple proposed infection dynamics into a single model while quantifying the uncertainty surrounding those effects, which also permits us to compare and contrast different approaches for modeling certain dynamics. The second advantage is the relative ease of interpretation of the effects estimated in our model, as well as the ease of interpretation of discrete-time hazard functions as the probability of a new infection for an uninfected farm. The third advantage is we are able to incorporate the infection status of source farms into our framework to improve our model fit.

## 2 Materials and methods

### 2.1 Study data

This study relies on several sources of swine production and disease occurrence data which have been described in greater detail elsewhere (Galvis, Corzo, Prada, et al. 2022b; Galvis, Jones, et al. 2022). Briefly, there are two main sources of data that we utilize: i) PEDV outbreak data, which provides records of new infections at the farm level through time; and ii) five sets of network data including animal movements and four different transportation vehicles, which are used to construct between-farm contact networks. We describe these data further in the following subsections.

#### 2.1.1 Farm outbreak data

Our outbreak data comes from three pig-producing systems with main operations in the United States. Due to confidentiality agreements, we will refer to production systems A, B, and C. The farms recorded the day each farm had a PEDV outbreak, and the farms are identified with a national premise identification (PremID). This PremID uniquely identifies a farm between multiple data sources and allows matching the outbreak data with the contact network data which includes animal and vehicle movements. The three production systems voluntarily report outbreaks to the Morrison Swine Health Monitoring Project (MSHMP), a Swine Health Information Center (SHIC) funded project and the source of the outbreak data.

The PEDV outbreak data we use to fit our model begins in January 2018 and extends through December 2020. During this time, there are 398 new PEDV outbreaks reported. We exclude 17 outbreaks that occur within 14 days of an earlier infection because we assume that these are the same event and the source of those rebreaks is not external, and we have developed a farm-level spread model. Here we consider rebreaks as in Galvis, Corzo, Prada, et al. 2022b, where they are considered to be a new infection caused by continued viral shedding of pigs or to be a result of inadequate cleaning and disinfection. We also include nine new infections in January 2021 as out-of-sample testing data. Real-world logistics affect the date the infection is recorded, and there is a delay in getting a positive diagnosis reported timely. Given these delay dynamics, we model the outbreak data at the weekly level.

Each farm is classified into a farm type based on the type of animals housed and raised. Typically, each site can be classified as a sow, nursery, finisher, or other production site (Galvis, Jones, et al. 2022). Generally, sow farms house breeding-age animals and suckling piglets, nurseries are where recently weaned pigs are housed for six-nine weeks before being transferred to finisher farms, and then pigs remain at finisher farms for 16-18 weeks before going to market (*Swine Industry Manual* 2011). A site may also be a gilt development unit (GDU) where selected females are raised to then be introduced to sow farms as replacement animals. It is possible that a single farm has multiple farm types, but for this study, it is considered a unique farm.

Furthermore, each farm has a primary accredited veterinarian who is an employee contracted by the farm’s swine company. The number of veterinarians employed varies by system, and veterinarians serve specific but potentially large geographic regions as well as a significant number of farms. They may be specialized in certain types of farms within a system and/or within a region (for example, only breeding farms).

The farms used in our model tend to be clustered such that there are concerns of local transmission of disease. This local transmission includes the propagation of PEDV among different systems (Galvis, Corzo, Prada, et al. 2022b; Machado et al. 2019).

#### 2.1.2 Network data

In addition to outbreak data, each of the three systems provided movements of pigs within their systems covering the period from November 29, 2019 until June 25, 2021. This data consists of the movement of weaned pigs from sow farms to nurseries and then from nurseries to finishers, though there are exceptions (Passafaro et al. 2020). Each recorded movement consists of a source PremID, a destination PremID, the date of the movement, and the total number of pigs moved. There is a significant amount of literature demonstrating that incoming pig movements increase the risk of disease spread (Galvis, Corzo, Prada, et al. 2022b; Galvis, Jones, et al. 2022; Passafaro et al. 2020; VanderWaal et al. 2018).

In addition to the pig-movement data, we have four vehicle-related networks. These four networks are analyzed in (Galvis, Corzo, and Machado 2022), and we rely on their processing and analysis of the data to incorporate the data into our model. The four additional networks are used to model the indirect contact by vehicles coming into farms. The networks correspond to the vehicles used for feed, animal delivery to farms, animal delivery to markets, and the personnel (crew) involved in the loading and unloading of pigs (Galvis, Corzo, and Machado 2022). We refer to these networks as the feed, pig, market, and labor vehicle networks, respectively.

It has been demonstrated that PEDV remains viable on a vehicle as it moves between farms (Wu et al. 2022), increasing the risk of spreading the disease to the destination farm if the source farm is infected (Büttner and Krieter 2022; Dee, Torremorell, et al. 2007; Dee, Deen, et al. 2004; Galvis, Corzo, and Machado 2022). Because of this risk, vehicles are typically cleaned and disinfected between farm visits (Dee, Torremorell, et al. 2007; Dee, Deen, et al. 2004). The data analyzed in (Galvis, Corzo, and Machado 2022; Galvis, Corzo, Prada, et al. 2022a) includes the times when the vehicles are cleaned and disinfected.

It is assumed that when a vehicle is sanitized it is completely successful, nullifying any chance of spreading an infection to the next visited farm. Therefore, our data for these four networks consists of the movements of the vehicles from a national PremID to another national PremID without the vehicle being sanitized in between. Each such observation includes a date as well as the time spent at the destination farm, which was computed in (Galvis, Corzo, and Machado 2022) by using the GPS data from the vehicles themselves. Data on 159 feed trucks, 118 pig trucks, 89 market trucks, and 32 crew trucks were used to construct contact networks, and a farm visit was defined by a vehicle remaining stationary for at least five minutes within 1.5 km of a farm. See (Galvis, Corzo, and Machado 2022) Section 2.1 for additional details.

### 2.2 Statistical analysis

We use time-to-event methods, also known as survival analysis (Klein and Moeschberger 2003; Klein, van Houwelingen, et al. 2013; Tutz and Schmid 2016). These methods allow modeling the factors that lead to certain farms becoming infected sooner than other farms (Klein and Moeschberger 2003; Klein, van Houwelingen, et al. 2013; Tutz and Schmid 2016). Because our data are weekly and farms can have multiple infection periods, we use a discrete-time, recurrent-event model (Tutz and Schmid 2016; Willett and Singer 1995). The infection status is defined as *Y* (*f,t*) = 1 if farm *f* is infected with PEDV in week *t* and *Y*(*f, t*) = 0 otherwise. The hazard function is the probability of a new infection at time t given the farm is not infected at time *t* – 1 and the history through time *t*:

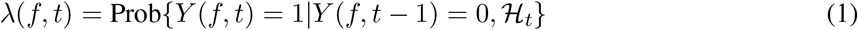

where the history 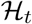 includes the covariates up to and including *t* as well as the infection status of all farms prior to *t*. We model the hazard probabilities using logistic regression (McCullagh and Nelder 1989; Tutz and Schmid 2016):

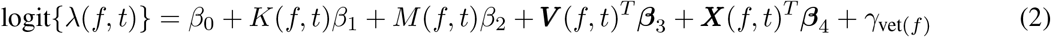

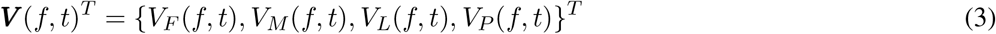

where *K* (*f,t*) is a local-transmission effect; *M* (*f,t*) is a movement-network effect; ***V*** (*f,t*) is the vector of the feed, market, labor, and pig vehicle-network effects; ***X***(*f,t*) is a vector of seasonality effects and other covariates; *γ*_vet_(*f*) is a random effect corresponding to the veterinarian of farm *f*; and 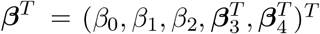 is a vector of fixed effects. Bold notation is used to indicate vectors.

The PEDV outbreak data is transformed from daily outbreaks into being weekly, but the transmission dynamics occur on a sub-weekly scale. Our solution to allow for sub-weekly dynamics is to introduce latent (i.e., not observed) intermediate time steps. Our final model for PEDV infections is

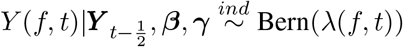

where *λ*(*f, t*) depends on the latent time steps. Details on the values of *Y*(*f, t*) at the latent time steps and the full Bayesian model specification are provided in Supplementary Sections 1.1 and 1.3, respectively.

Our formulations for *K*(*f,t*), *M*(*f,t*), and ***V***(*f,t*) depend on interactions with the PEDV infection status of farms besides *f* itself. However, the PEDV infection data only includes the times a new infection is observed for each farm, not for how long the farm stayed infected. The length of time a pig-producing farm is infected is understood to vary by farm type (Goede and Morrison 2016). We adopt the assumptions made in (Galvis, Jones, et al. 2022) and assume that nurseries have an infection length of seven weeks, GDUs take 22 weeks, finishers take 25 weeks, and sow farms take 28 weeks (Sanhueza, Vilalta, et al. 2019). In the following sections, we will provide overviews of our formulations for these effects as well as our model-selection criteria to choose between them.

#### 2.2.1 Network effects

We form a directed graph based on the movement data described in Section 2.1.2, where the edges and directions are defined using the observed movements. We compute *M* (*f,t*) in (2) as a summary statistic of the interaction of this directed graph and the infection statuses of the farms in the network. Let 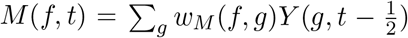, where *w_M_*(*f, g*) is the edge weight on the directed graph from farm *g* to farm *f*. We consider two formulations for *w_M_*(*f,g*). The first is to let *w_M_*(*f,g*) = 1 if farm *g* ever sent pigs to farm f and let it equal 0 otherwise. The second formulation is to set *w_M_* (*f, g*) equal to the number of animals shipped from *g* to *f*, which gives more weights to edges with more animals being moved. To decide between these two formulations, we use the model-selection criteria discussed in Section 2.2.4.

To model the indirect contacts between farms using the data for the four vehicle networks outlined in Section 2.1.2, we proceed in a similar fashion to *M*(*f, t*). We construct a directed graph based on the movements of the vehicles, where *w_F_* (*f,g*) = 1 if a directed edge exists for the feed network from farm *g* to farm *f*. Similar weights are defined analogously for the pig, market, and labor vehicle networks. Then 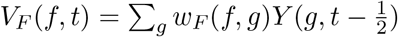, with *V_p_*(*f, t*), *V_M_*(*f, t*), and *V_L_*(*f, t*) being defined in the same manner.

We consider an alternative approach for the vehicle-network effects. Because the vehicles visit many farms in sequence, we consider calculating the elements of ***V***(*f, t*) by summarizing the time spent at *f* by trucks arriving after having visited an infected farm since the last disinfection. Details of this approach are provided in Supplementary Section 1.2. Again, to decide between these two formulations, we use the model-selection criteria discussed in a later Section 2.2.4.

#### 2.2.2 Local-infection effects

It is widely accepted that a major source of PEDV infections can disseminate from farm to farm given their spatial proximity (Galvis, Corzo, Prada, et al. 2022b; Galvis, Jones, et al. 2022; VanderWaal et al. 2018). Previous studies have suggested PEDV can survive for up to 20 days outside a host (Kim et al. 2018) and is able to propagate at least 16 km by wind (Alonso et al. 2014), and therefore we expect that a farm’s hazard function increases as a result of being near other farms (Galvis, Corzo, Prada, et al. 2022b). A widely adopted approach to modeling local transmission is through transmission kernels, which we use in our modeling framework. These mechanistic models typically model spread as a function of infection status, farm populations, and distances between farms (Flood et al. 2013; Tildesley et al. 2012). A common local-infection kernel used to model local transmission is the gravity model (Gog et al. 2014; Viboud et al. 2006; Xia et al. 2004), which is typically written as:

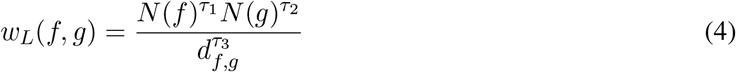

where *N* (*f*) is the population of farm *f, N* (*g*) is the population of farm *g*, and *d_f,g_* is the distance between farm *f* and *g*. We follow (Galvis, Corzo, Prada, et al. 2022b; Galvis, Jones, et al. 2022) and assume *τ*_1_ = *τ*_2_ = 1 and *τ*_3_ = 2.

Some recent work suggests the gravity model is not ideal to model transmission because it does not account for “intervening” farms (Bjørnstad et al. 2019). Specifically, the effect of *g* on f is expected to be diminished if a third farm *h* is between f and *g,* which cannot be captured by the gravity model. A proposed alternative to the gravity model that accounts for intervening farms is the Stouffer model (Bjørnstad et al. 2019). Assuming all animals in the farm are either infected or not, the Stouffer model is written as:

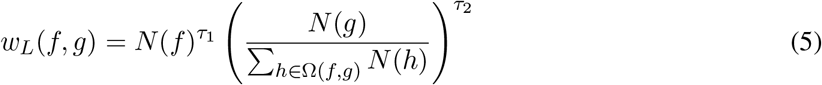

where Ω(*f, g*) is the set of farms whose distance from f is no more than the distance between *f* and *g*. We assume *τ*_1_ = *τ*_2_ = 1 to be consistent with the gravity-model formulation.

We interact our two proposed kernels with the farm-infection data to calculate *K*(*f, t*). In both cases, we only consider farms that are separated by at most 30 km so that *w_L_*(*f, g*) = 0 if > 30 km, consistent with Galvis, Jones, et al. 2022. Furthermore, we found that because these local-transmission effects could be numerically large, taking the natural logarithm of the effects improved model fit. Therefore K(f, t) may be calculated as:

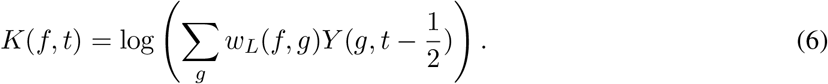

In the case of no neighboring infected farms in week *t*, we define *K*(*f, t*) = 0. We decide between the gravity and Stouffer formulations using the model-selection criteria in 2.2.4.

#### 2.2.3 Additional covariates and seasonality effects

In addition to the network and local-infection effects, there are additional covariates that impact a farm’s hazard function. The most well-known of these effects is the seasonality of PEDV spread (Goede and Morrison 2016). Specifically, PEDV tends to spread more easily in the winter than in the summer because the virus is able to survive longer without a host (Lee 2015; Machado et al. 2019). Therefore, we incorporate seasonality effects into our model through the use of monthly dummy variables. Another potentially impactful covariate is how much vegetation there is near a farm, which we measure using the Enhanced Vegetation Index (EVI) (Vermote et al. 2016). Some studies have suggested that vegetation coverage may help prevent the local transmission of PEDV between farms (Arruda et al. 2017; Galvis, Corzo, Prada, et al. 2022b; Jara et al. 2021), and the effect might be stronger in winter when PEDV can survive longer without a host (Lee 2015; Machado et al. 2019). The EVI data are reported every 16 days, and we calculate a simple weighted average of EVIs if a week *t* overlaps two EVI reporting periods. Finally, we include the elevation in meters of the farms because some studies have suggested that elevation could have an effect in the propagation of swine diseases (Arruda et al. 2017; Jara et al. 2021; Sanhueza, Stevenson, et al. 2020).

Survival models allow random effects (frailties) that account for variability in the subjects being studied (Klein and Moeschberger 2003; Tutz and Schmid 2016). These random effects allow survival curves to vary by individual even after controlling for fixed effects. In the case of our model, after accounting for the farm-level covariates, networks, and local transmission, individual farms have additional unobserved features that lead to innately higher or lower hazard functions. However, many farms do not have any observed outbreaks during the time periods considered, leading to difficulties in estimating farm-level random effects. Our solution is to use independent random effects for veterinarians. Following from the description in Section 2.1.1, using veterinarians as random effects works as a proxy for the interaction of system, farm type, and spatial region.

#### 2.2.4 Model selection

We have proposed different network formulations in Section 2.2.1 and also two local-infection formulations in Section 2.2.2. To decide between the different model specifications, we use the Watanabe-Akaike information criteria (WAIC), a well-known criterion that approximates leave-one-out cross-validation in Bayesian models (Reich and Ghosh 2019). Lower WAICs are better, and a decrease in ten or more in WAIC is considered to be a loose rule of thumb for an improvement in model fit (Reich and Ghosh 2019). We use the WAICs to evaluate in-sample fit of our proposed formulations.

We also consider out-of-sample performance using the January 2021 infection data. To assess predictive performance, we rely on two measures. The first is the well-known area under the receiver-operating-characteristic curve (AUC), which measures the performance of a classifier based on its true-positive rate and its false-positive rate (Saito and Rehmsmeier 2015). The area under the precision-recall curve (PR AUC) may be better at analyzing the performance of a binary classifier for data with a large proportion of zeroes (Saito and Rehmsmeier 2015). The PR AUC measures the trade-off between the true-positive rate and the classifier’s precision. The baseline performance for a binary classifier measured by AUC is 0.5, meaning that any binary classifier should have an AUC better than 0.5 (Saito and Rehmsmeier 2015). The baseline performance for a binary classifier, when measured by PR AUC, is the percentage of positives in the test data (Saito and Rehmsmeier 2015). For our analysis, the PR AUC baseline is 0.001. Our calculations for AUC and PR AUC are for all four weeks in January 2021 simultaneously and are calculated after estimating the latent time steps. Additional details are in Supplementary Section 1.5.

## 3 Results

Our proposed model was fit to the PEDV infection data from 2018 to 2020. We present our results in three sections. In the first section, we show our results for the different formulations of the network and local effects. In the second section, we compare our chosen model with a simpler model that does not incorporate infection status into the effects. In the third section, we present the point estimates and credible intervals for our final model. In the final section, we present analytic results using the final model.

### 3.1 Model selection for local-transmission and network effects

We present our results for fitting our model using the different formulations for the movement, vehicle, and local effects. Briefly, we evaluate weighting the directed edges of the movement network by animals “animals” or not “binary”, whether it is better to use a directed graph for the vehicle networks “binary” or to use time spent visiting farms “time”, and whether the gravity or Stouffer kernel should be used for local-transmission effects. Table 1 below presents the main findings.

**Table 1:**
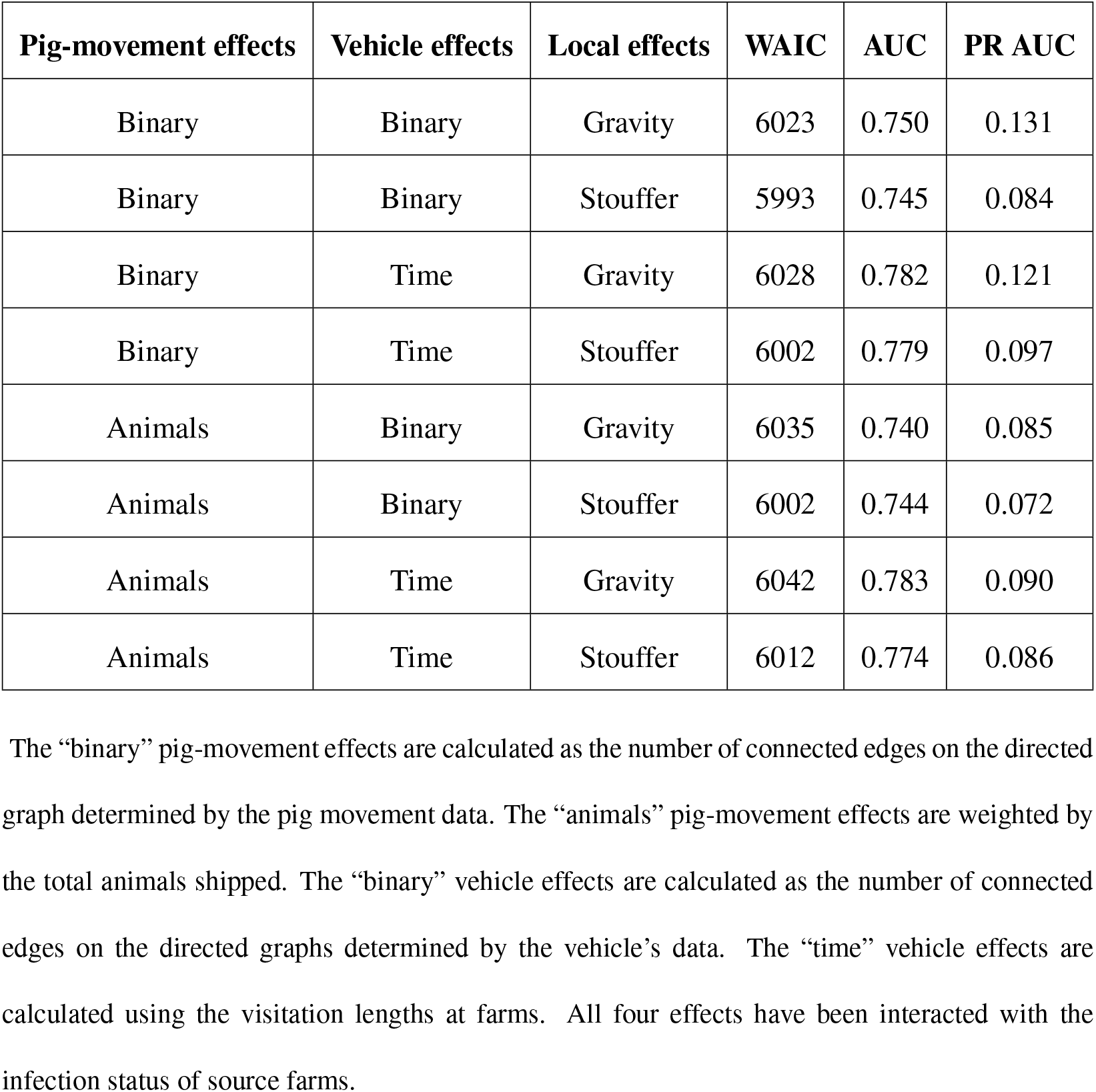
Model selection for different proposed network and local-transmission effects.

The “binary” pig-movement effects are calculated as the number of connected edges on the directed graph determined by the pig movement data. The “animals” pig-movement effects are weighted by the total animals shipped. The “binary” vehicle effects are calculated as the number of connected edges on the directed graphs determined by the vehicle’s data. The “time” vehicle effects are calculated using the visitation lengths at farms. All four effects have been interacted with the infection status of source farms.

Our results in Table 1 show several sound choices for best model. Measured by WAIC, the Stouffer formulation for the local effects fits the in-sample data better than the gravity formulation. By that measure, the binary pig-movement effects, the binary vehicle effects, and the Stouffer local effects yields the best model. However, as described in Section 2.2.4 an increase of nine for the WAIC is not typically considered to be a material difference in model fit (Reich and Ghosh 2019), suggesting the two models with 6002 WAIC may fit the in-sample data just as well. In such a case, we prefer the formulation using binary pig-movement effects and time vehicle effects, taking the AUC and PR AUC into consideration.

However, the best out-of-sample predictions when measured by PR AUC are associated with the binary pig-movement effects, the binary vehicle effects, and the gravity model for local transmission. Our work has consistently shown worse WAICs for the gravity model, so we still prefer the Stouffer formulation.

We choose to use the model with formulations of binary pig-movement effects, time vehicle effects, and Stouffer local effects. This model has the strongest overall performance using the three criteria in Table 1. Nevertheless, some readers may prefer either of the models that use the binary pig-movements effects or binary vehicle effects by putting preferential weight on WAIC or PR AUC. We provide those results in Supplementary Section 2.

### 3.2 The importance of interacting network and local effects with infections

Our model formulation is driven by the inclusion of *Y*(*f, t*) in the network and local-transmission effects. Its inclusion leads to a more complex model, but we have found consistently that the additional complexity is justified.

To demonstrate the need for including the infections in the network and local-transmission effects, we compare our chosen with a comparable, simpler mixed-effects logistic regression with the same structure as our model presented in (2) but make the following changes:

- The effects *M* (*f,t*), *K* (*f,t*), and ***V*** (*f,t*) do not depend on *Y* (*f, t*) as described in Sections 2.2.1 and 2.2.2.
- Because the fixed effects no longer depend on the responses *Y*(*f, t*), we do not incorporate the latent time steps to make it a valid statistical model.

Because the simpler, mixed-effects model will have fewer observations because of the omission of the latent time steps, a comparison made by WAIC is not straightforward (the textbook (Reich and Ghosh 2019) provides an in-depth overview of WAIC in model selection). Instead, we compare out-of-sample performance between our chosen model from Section 3.1 and an otherwise identical model with the two indicated adjustments above. As result, we have consistently found similar results in our work to those in Table 2. Our results suggest that tracking infected farms is important to better model the dynamics of the spread of PEDV (Table 2).

**Table 2:**
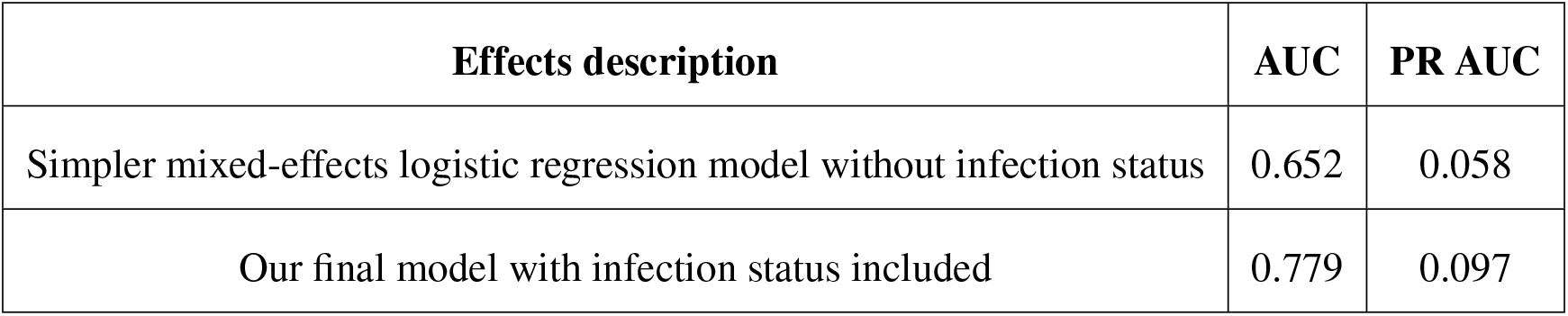
Comparison of models with and without inclusion of infection status in the network and localinfection effects.

### 3.3 Summary of chosen final model

Table 3 summarizes the point estimates for the fixed effects and provides their 95% credible intervals. For the monthly dummies representing the seasonality effects, July is omitted as a baseline month.

**Table 3:**
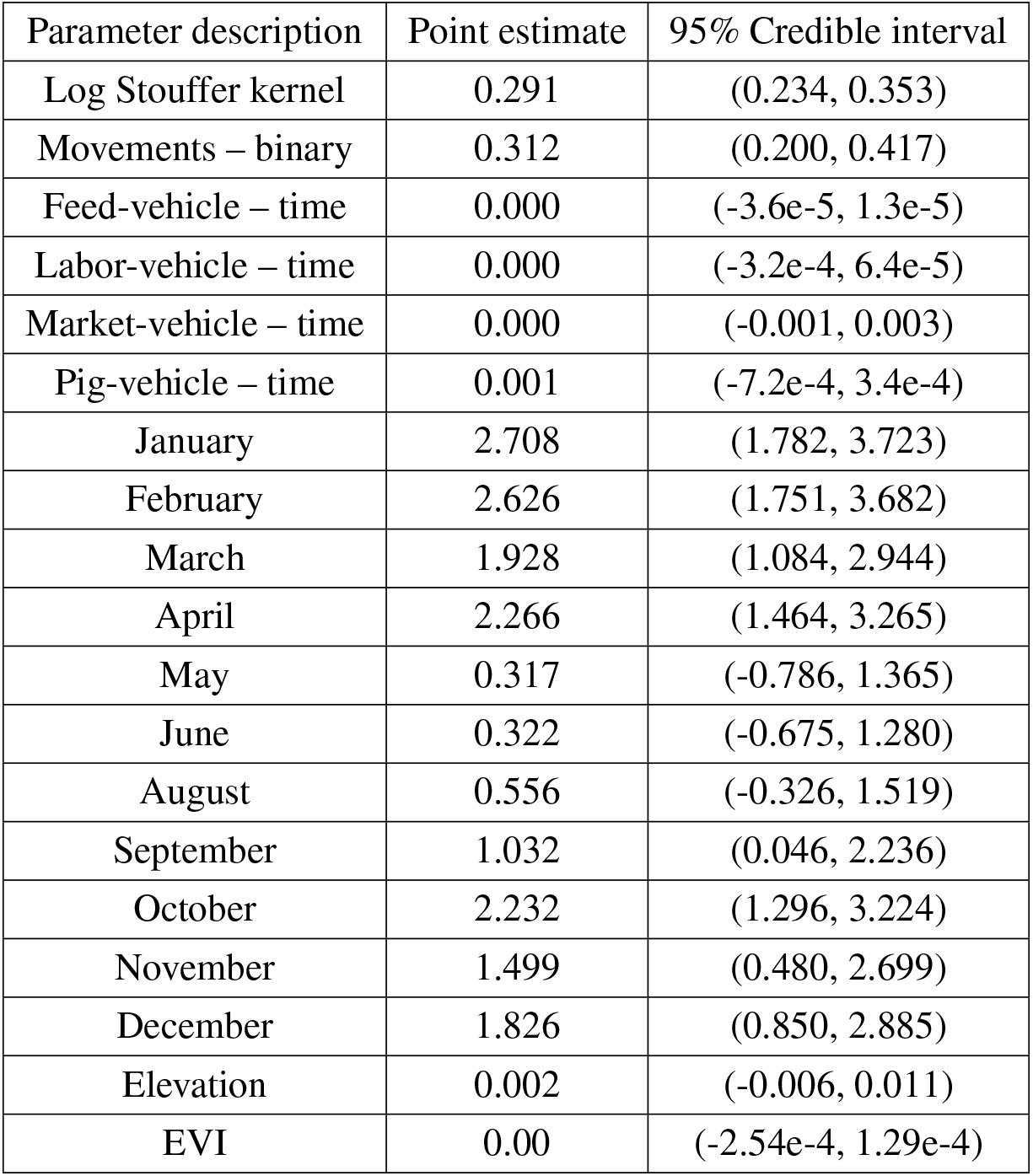
PEDV model fixed-effects estimates.

Our results show there is significant evidence there are local-transmission effects, pig-movement effects, and seasonality effects affecting between-farm transmission of PEDV. These are evidenced by the positive coefficients and the 95% credible intervals not containing zero for the log Stouffer transmission kernel, the binary pig-movements network, and the months October-April. The 95% credible intervals for May, June, August, and September all include zero, suggesting that the seasonality effects of those months are not significantly different than the baseline effect of July.

The credible intervals for all four vehicle-network coefficients as well as EVI and farm elevation included zero. The EVI and elevation do not vary dramatically over our study region, so having a larger study area that includes more variability in elevation and greenery may allow these effects to be better estimated.

### 3.4 Analysis and interpretation

Previous studies have proposed models for PEDV infections that have used grid-based predictive performance in lieu of farm-level predictive performance (Galvis, Jones, et al. 2022; Machado et al. 2019). These grid-based predictions are useful to assess how well the model can differentiate regions where the infection is likely to occur from regions where new infections are less likely. In this study, we calculate AUC and PR AUC for three different grid sizes: a 10 km x 10 km grid, a 5 km x 5 km grid, and a 1 km x 1 km grid. We assess predictive performance in grid cells where there is at least one farm. Table 4 summarizes our results.

**Table 4:**
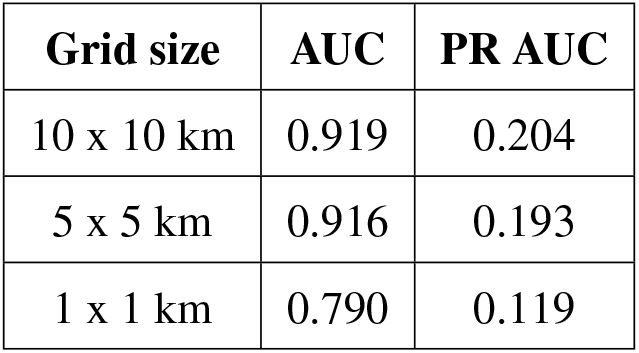
Summary of grid-based AUCs and PR AUCs.

These results suggest our model can perform very well in identifying local areas where PEDV outbreaks are more likely to occur, which allows stakeholders to make more informed decisions on which farms should be prioritized for control measures. In particular, the results using a 5 km × 5 km grid are nearly as good as a much larger 10 km × 10 km grid.

We next turn to describing the interpretation of our model presented in Table 3. As an illustrative example, consider the seasonality hazard functions plotted in Figure 1. These hazard functions are calculated using typical values for the effects while varying only the month of the year. The black line shows the typical probabilities of a new infection for an uninfected farm during a time step in the specified month, while the red lines show the probabilities but using the 2.5% and 97.5% quantiles for the seasonality effects. As Figure 1 shows, there is a significant amount of uncertainty in these seasonality effects, even though there is evidence the colder months have a higher probability of a new PEDV infection.

**Figure 1.**
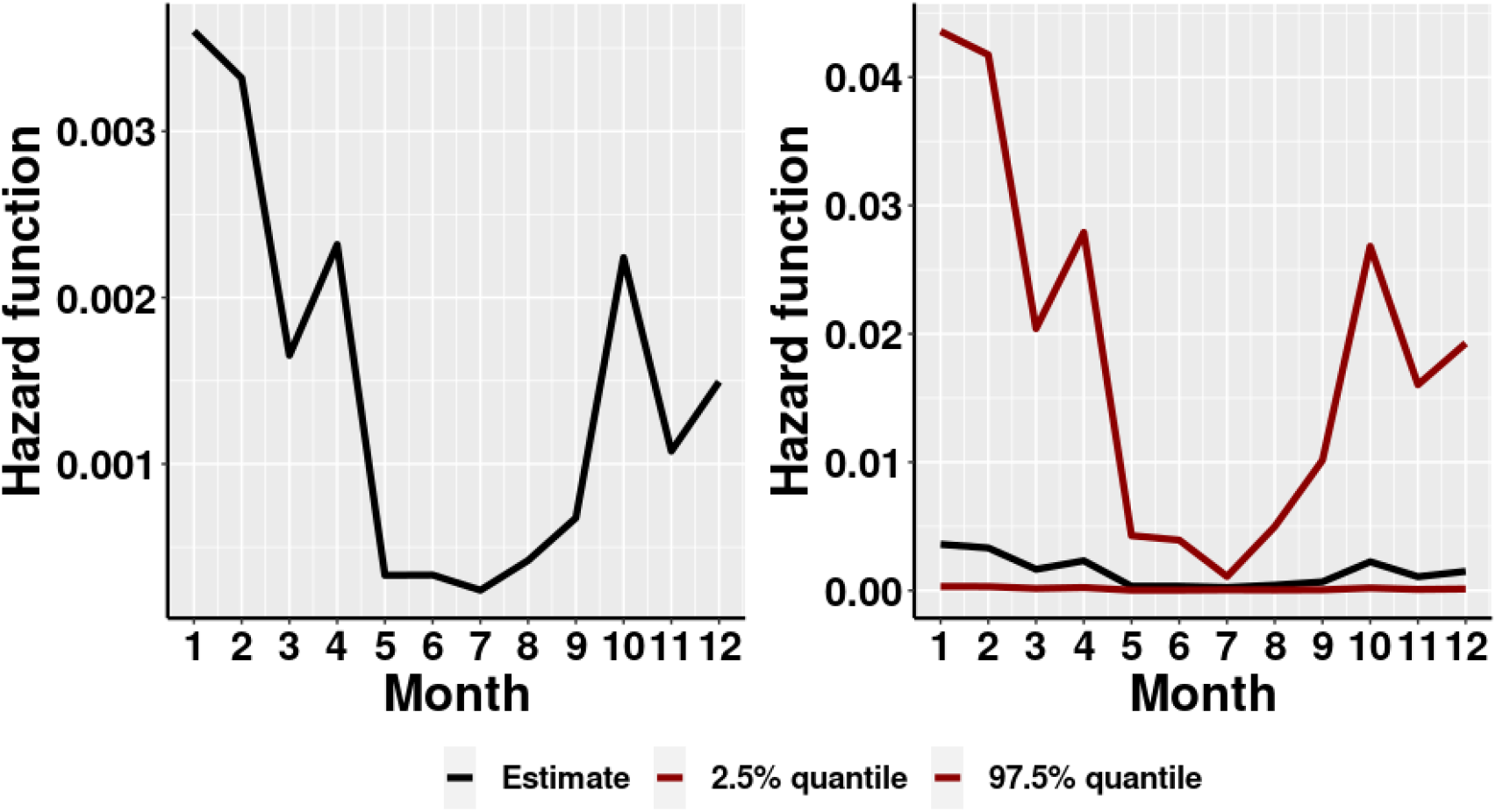
Typical seasonality hazard function (left) with 95% credible interval (right, scale changed)

The log of the Stouffer local-transmission kernel is difficult to interpret. It is a function of the size of nearby farms, the distance from farm to farm, and if those farms are infected. To help interpret this effect, in Figure 2 we calculate 1 – the estimated survival function for example sets of farm characteristics (starting in January) but varying the log Stouffer kernel to “low”, ‘‘medium”, and “high” values based on the data used in our model. The low kernel corresponds to a farm that was 15 km from the closest infected farm but with many uninfected farms closer than 15 km. The medium kernel corresponds to a farm that is 2 km from the closest infected farm. The high kernel corresponds to a farm that is 5 km from the closest infected farm, but there are more than 20 infected farms within 30 km. In our model, *K*(*f, t*) varies by week, but we hold it fixed for these example values below for illustration. There is a complex local-spread dynamic where both distance and the number of infected farms play a part.

**Figure 2:**
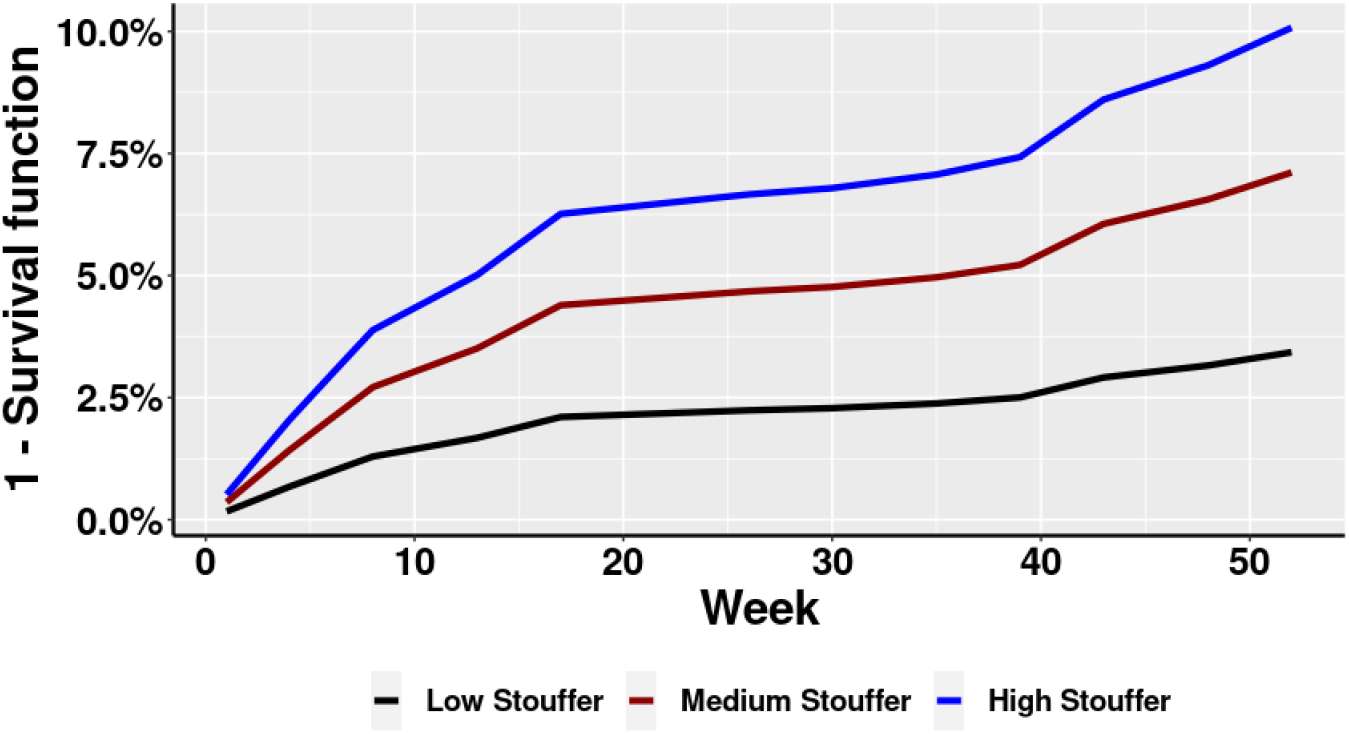
Survival functions for low, median, and high values of the log Stouffer transmission kernel.

## 4 Discussion

In this paper we have proposed a model that achieved two competing goals: a flexible modeling framework that allows capturing the spread dynamics of PEDV, and our model can be fit on any modern computer in a reasonable runtime. The benefit of achieving these two goals is that we were able to evaluate many different proposed effects formulations, and the success of our approach is evidenced by the quality of our out-of-sample predictions at both the farm level and the grid level. Our work can be readily generalized to modeling the farm-level spread of other diseases, and its simplicity means it can rapidly fit data on new and emerging pathogens. A key component of our modeling framework is the inclusion of farm-infection status in the calculations of the local-transmission and network effects. We found this inclusion consistently improved the fit of our models no matter the criterion used. Our findings imply that what matters is not being near other farms but being near other infected farms; likewise, what matters is not receiving pigs from another farm but receiving infected pigs. Thus, our results demonstrated that both effects should be considered.

Our proposed model is a valid statistical model, which is not a trivial conclusion given the context of our framework. Though this concern is technical, it is worth emphasizing. Because our data is weekly, implementing a simpler version of our model such that *Y*(*f, t*) depends on the other farms at the same time point *t* is not necessarily a valid model. Here “valid” means our conditionally specified model can be used to calculate the joint probability of multiple farms becoming infected in the same week. The interested reader should refer to Joe and Liu 1996 for further details. Because of our introduction of the latent time steps and conditioning values at time *t* on 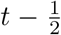, our model is valid. Therefore, the key to our methodology is including farm-infection status in the model effects, and we have also provided a solution to the technical complication that follows from this inclusion.

There have been many proposed dynamics for the spread of PEDV, such as local transmission (Alonso et al. 2014; Alvarez et al. 2016; Beam et al. 2015) and indirect contacts from vehicles (Büttner and Krieter 2022; Dee, Torremorell, et al. 2007; Dee, Deen, et al. 2004; Galvis, Corzo, and Machado 2022). We considered these different dynamics in a single model. Because of our access to private movement and vehicle data, we were able to model network effects while still controlling for local transmission, seasonality, EVI, and elevation. If these dynamics influence PEDV transmission, their omission can lead to biased parameter estimates. This omitted-variable bias has been discussed in other fields as impacting regression models (Katicha and Flintsch 2022; Rinella et al. 2020; Stevenson 2018). For example, excluding network effects can lead to the coefficient on the local-transmission effects to increase greatly in magnitude. Therefore, comprehensive models such as ours help to avoid these concerns.

We have made a few assumptions in our model. The first is we used static networks for the pig movements and vehicle networks. However, recent work has suggested these networks have high causal fidelity, meaning the static networks approximate the temporal networks well (Galvis, Corzo, and Machado 2022). Likewise, we have assumed that the 2018 and 2019 networks were the same as the 2020 networks, which we believe is reasonable given the aforementioned high causal fidelity. We only have vehicle-movement data for one pig-producing company, which may mean we are under-representing the true indirect contacts between farms. Nevertheless, our model can only be improved by incorporating additional vehicle data, and it may allow us to better estimate the effect of indirect contacts between farms. Additional infection data may also allow us to estimate the parameters in the gravity and Stouffer local-transmission kernels. Also, daily infection data that captures the true day of a new infection would not require the latent time steps, and it would remove technical concerns about valid joint models.

## 5 Conclusion

We fit a model to a comprehensive PEDV dataset consisting of three years of farm-level infections. Our model estimated the effects of different dynamics proposed in the literature as affecting the spread of PEDV, including local transmission, the movement of pigs, and the spread of disease by vehicles visiting farms. Our results show a significant effect of local transmission and pig movements, and we have demonstrated the importance of accounting for the infection status of other farms when fitting a PEDV model.

## Acknowledgments

Authors would like to acknowledge participating companies and veterinarians.

## Authors’ contributions

PT, BJR, and GM conceived the study. CC coordinated the disease data collection through the Morrison Swine Health Monitoring Project (MSHMP) system. PT conducted data processing and cleaning, designed the model, and wrote the model-fitting code. PT designed the computational analysis. PT, BJR, and GM wrote and edited the manuscript. All authors discussed the results and critically reviewed the manuscript. GM and BJR secured the funding.

## Conflict of interest

All authors confirm that there are no conflicts of interest to declare

## Ethical statement

The authors confirm the ethical policies of the journal, as noted on the journal’s author guidelines page. Since this work did not involve animal sampling nor questionnaire data collection by the researchers, there was no need for ethics permits.

## Data Availability Statement

The data that support the findings of this study are not publicly available and are protected by confidential agreements, therefore, are not available.

## Funding

This work was supported by Food and Agriculture Cyberinformatics and Tools, 2020-67021-32462 from the USDA National Institute of Food and Agriculture. The Morrison Swine Health Monitoring Project is a Swine Health Information Center (SHIC) funded project.

## Notes

### Competing Interest Statement

The authors have declared no competing interest.

