## Supplementary info for "A discrete-time survival model for porcine epidemic diarrhea virus"

$$\lambda(f,t) = \text{Prob}\{Y(f,t) = 1 | Y(f,t-1) = 0, \mathcal{H}_t\} \quad (1)$$

where the history $\mathcal{H}_t$ includes the covariates up to and including $t$ as well as the infection status of all farms prior to $t$. We model the hazard probabilities using logistic regression (McCullagh and Nelder 1989; Tutz and Schmid 2016):

$$\text{logit}\{\lambda(f,t)\} = \beta_0 + K(f,t)\beta_1 + M(f,t)\beta_2 + \boldsymbol{V}(f,t)^T\boldsymbol{\beta}_3 + \boldsymbol{X}(f,t)^T\boldsymbol{\beta}_4 + \gamma_{\text{vet}(f)} \tag{2}$$

$$\boldsymbol{V}(f,t)^T = \{V_F(f,t), V_M(f,t), V_L(f,t), V_P(f,t)\}^T \tag{3}$$

where $K(f,t)$ is a local-transmission effect; $M(f,t)$ is a movement-network effect; $\boldsymbol{V}(f,t)$ is the vector of the feed, market, labor, and pig vehicle-network effects; $\boldsymbol{X}(f,t)$ is a vector of seasonality effects and other covariates; $\gamma_{\text{vet}(f)}$ is a random effect corresponding to the veterinarian of farm $f$; and $\boldsymbol{\beta}^T = (\beta_0, \beta_1, \beta_2, \boldsymbol{\beta}_3^T, \boldsymbol{\beta}_4^T)^T$ is a vector of fixed effects. Bold notation is used to indicate vectors.

The PEDV outbreak data is transformed from daily outbreaks into being weekly, but the transmission dynamics occur on a sub-weekly scale. Our solution to allow for sub-weekly dynamics is to introduce latent (i.e., not observed) intermediate time steps. Our final model for PEDV infections is

$$Y(f,t)|\boldsymbol{Y}_{t-\frac{1}{2}}, \boldsymbol{\beta}, \boldsymbol{\gamma} \overset{ind}{\sim} \text{Bern}(\lambda(f,t))$$

where $\lambda(f,t)$ depends on the latent time steps. Details on the values of $Y(f,t)$ at the latent time steps and the full Bayesian model specification are provided in Supplementary Sections 1.1 and 1.3, respectively.

$$w_L(f,g) = \frac{N(f)^{\tau_1} N(g)^{\tau_2}}{d_{f,g}^{\tau_3}} \tag{4}$$

where $N(f)$ is the population of farm $f$, $N(g)$ is the population of farm $g$, and $d_{f,g}$ is the distance between farm $f$ and $g$. We follow (Galvis, Corzo, Prada, et al. 2022b; Galvis, Jones, et al. 2022) and assume $\tau_1 = \tau_2 = 1$ and $\tau_3 = 2$.

Some recent work suggests the gravity model is not ideal to model transmission because it does not account for "intervening" farms (Bjørnstad et al. 2019). Specifically, the effect of $g$ on $f$ is expected to be diminished if a third farm $h$ is between $f$ and $g$, which cannot be captured by the gravity model. A proposed alternative to the gravity model that accounts for intervening farms is the Stouffer model (Bjørnstad et al.

2019). Assuming all animals in the farm are either infected or not, the Stouffer model is written as:

$$w_L(f,g) = N(f)^{\tau_1}\left(\frac{N(g)}{\sum_{h\in\Omega(f,g)} N(h)}\right)^{\tau_2} \tag{5}$$

where $\Omega(f, g)$ is the set of farms whose distance from $f$ is no more than the distance between $f$ and $g$. We assume $\tau_1 = \tau_2 = 1$ to be consistent with the gravity-model formulation.

We interact our two proposed kernels with the farm-infection data to calculate $K(f, t)$. In both cases, we only consider farms that are separated by at most 30 km so that $w_L(f, g) = 0$ if $d_{f,g} > 30$ km, consistent with Galvis, Jones, et al. 2022. Furthermore, we found that because these local-transmission effects could be numerically large, taking the natural logarithm of the effects improved model fit. Therefore $K(f, t)$ may be calculated as:

$$K(f,t) = \log\left(\sum_g w_L(f,g)Y(g,t-\frac{1}{2})\right). \tag{6}$$

In the case of no neighboring infected farms in week $t$, we define $K(f, t) = 0$. We decide between the gravity and Stouffer formulations using the model-selection criteria in 2.2.4.

Table 1: Model selection for different proposed network and local-transmission effects

| **Pig-movement effects** | **Vehicle effects** | **Local effects** | **WAIC** | **AUC** | **PR AUC** |
|---|---|---|---|---|---|
| Binary | Binary | Gravity | 6023 | 0.750 | 0.131 |
| Binary | Binary | Stouffer | 5993 | 0.745 | 0.084 |
| Binary | Time | Gravity | 6028 | 0.782 | 0.121 |
| Binary | Time | Stouffer | 6002 | 0.779 | 0.097 |
| Animals | Binary | Gravity | 6035 | 0.740 | 0.085 |
| Animals | Binary | Stouffer | 6002 | 0.744 | 0.072 |
| Animals | Time | Gravity | 6042 | 0.783 | 0.090 |
| Animals | Time | Stouffer | 6012 | 0.774 | 0.086 |

Table 2: Comparison of models with and without inclusion of infection status in the network and local-infection effects

| **Effects description** | **AUC** | **PR AUC** |
|---|---|---|
| Simpler mixed-effects logistic regression model without infection status | 0.652 | 0.058 |
| Our final model with infection status included | 0.779 | 0.097 |

##### 3.3 Summary of chosen final model

Table 3 summarizes the point estimates for the fixed effects and provides their 95% credible intervals. For the monthly dummies representing the seasonality effects, July is omitted as a baseline month.

Table 3: PEDV model fixed-effects estimates

| Parameter description | Point estimate | 95% Credible interval |
|---|---|---|
| Log Stouffer kernel | 0.291 | (0.234, 0.353) |
| Movements – binary | 0.312 | (0.200, 0.417) |
| Feed-vehicle – time | 0.000 | (-3.6e-5, 1.3e-5) |
| Labor-vehicle – time | 0.000 | (-3.2e-4, 6.4e-5) |
| Market-vehicle – time | 0.000 | (-0.001, 0.003) |
| Pig-vehicle – time | 0.001 | (-7.2e-4, 3.4e-4) |
| January | 2.708 | (1.782, 3.723) |
| February | 2.626 | (1.751, 3.682) |
| March | 1.928 | (1.084, 2.944) |
| April | 2.266 | (1.464, 3.265) |
| May | 0.317 | (-0.786, 1.365) |
| June | 0.322 | (-0.675, 1.280) |
| August | 0.556 | (-0.326, 1.519) |
| September | 1.032 | (0.046, 2.236) |
| October | 2.232 | (1.296, 3.224) |
| November | 1.499 | (0.480, 2.699) |
| December | 1.826 | (0.850, 2.885) |
| Elevation | 0.002 | (-0.006, 0.011) |
| EVI | 0.00 | (-2.54e-4, 1.29e-4) |

Our results show there is significant evidence there are local-transmission effects, pig-movement effects, and seasonality effects affecting between-farm transmission of PEDV. These are evidenced by the positive coefficients and the 95% credible intervals not containing zero for the log Stouffer transmission kernel, the binary pig-movements network, and the months October-April. The 95% credible intervals for May, June, August, and September all include zero, suggesting that the seasonality effects of those months are not significantly different than the baseline effect of July.

The credible intervals for all four vehicle-network coefficients as well as EVI and farm elevation included zero. The EVI and elevation do not vary dramatically over our study region, so having a larger study area that includes more variability in elevation and greenery may allow these effects to be better estimated.

##### 3.4 Analysis and interpretation

Previous studies have proposed models for PEDV infections that have used grid-based predictive performance in lieu of farm-level predictive performance (Galvis, Jones, et al. 2022; Machado et al. 2019). These grid-based predictions are useful to assess how well the model can differentiate regions where the infection is likely to occur from regions where new infections are less likely. In this study, we calculate AUC and PR AUC for three different grid sizes: a 10 km x 10 km grid, a 5 km x 5 km grid, and a 1 km x 1 km grid. We assess predictive performance in grid cells where there is at least one farm. Table 4 summarizes our results.

Table 4: Summary of grid-based AUCs and PR AUCs

| Grid size | AUC | PR AUC |
|---|---|---|
| 10 x 10 km | 0.919 | 0.204 |
| 5 x 5 km | 0.916 | 0.193 |
| 1 x 1 km | 0.790 | 0.119 |

These results suggest our model can perform very well in identifying local areas where PEDV outbreaks are more likely to occur, which allows stakeholders to make more informed decisions on which farms should be prioritized for control measures. In particular, the results using a 5 km $\times$ 5 km grid are nearly as good as a much larger 10 km $\times$ 10 km grid.

We next turn to describing the interpretation of our model presented in Table 3. As an illustrative example, consider the seasonality hazard functions plotted in Figure 1. These hazard functions are calculated using typical values for the effects while varying only the month of the year. The black line shows the typical probabilities of a new infection for an uninfected farm during a time step in the specified month, while the red lines show the probabilities but using the 2.5% and 97.5% quantiles for the seasonality effects. As

Figure 1 shows, there is a significant amount of uncertainty in these seasonality effects, even though there is evidence the colder months have a higher probability of a new PEDV infection.

Figure 1: Typical seasonality hazard function (left) with 95% credible interval (right, scale changed)

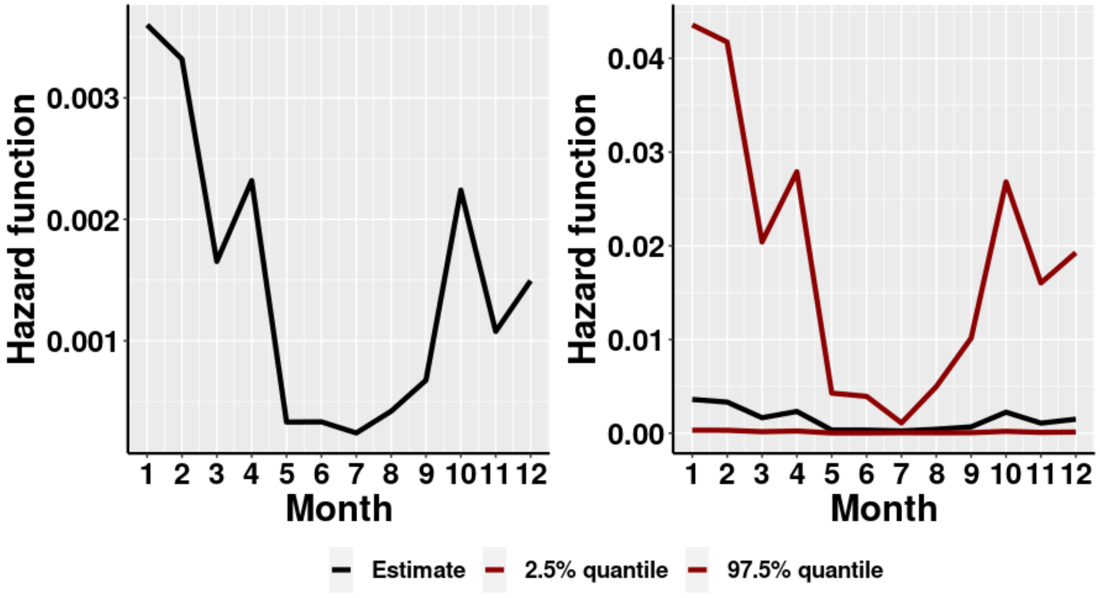


The log of the Stouffer local-transmission kernel is difficult to interpret. It is a function of the size of nearby farms, the distance from farm to farm, and if those farms are infected. To help interpret this effect, in Figure 2 we calculate 1 - the estimated survival function for example sets of farm characteristics (starting in January) but varying the log Stouffer kernel to ”low”, ”medium”, and ”high” values based on the data used in our model. The low kernel corresponds to a farm that was 15 km from the closest infected farm but with many uninfected farms closer than 15 km. The medium kernel corresponds to a farm that is 2 km from the closest infected farm. The high kernel corresponds to a farm that is 5 km from the closest infected farm, but there are more than 20 infected farms within 30 km. In our model, $K(f, t)$ varies by week, but we hold it fixed for these example values below for illustration. There is a complex local-spread dynamic where both distance and the number of infected farms play a part.

Figure 2: Survival functions for low, median, and high values of the log Stouffer transmission kernel

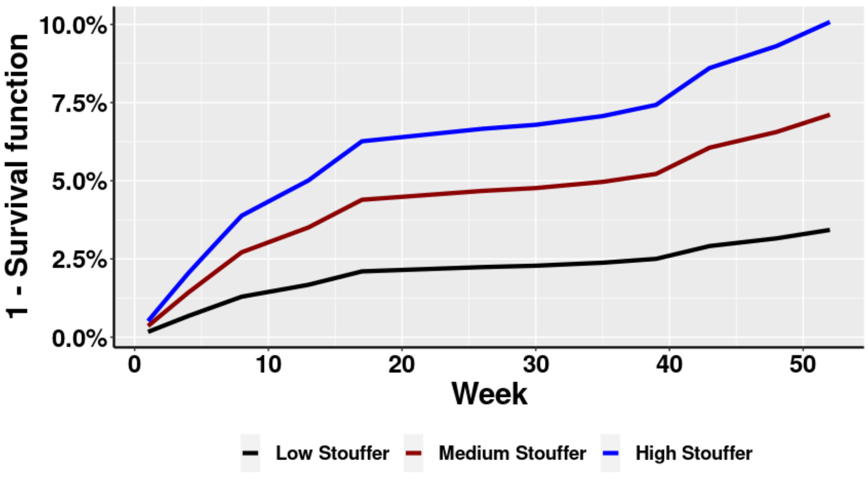


### 4 Discussion

In this paper we have proposed a model that achieved two competing goals: a flexible modeling framework that allows capturing the spread dynamics of PEDV, and our model can be fit on any modern computer in a reasonable runtime. The benefit of achieving these two goals is that we were able to evaluate many different proposed effects formulations, and the success of our approach is evidenced by the quality of our out-of-sample predictions at both the farm level and the grid level. Our work can be readily generalized to modeling the farm-level spread of other diseases, and its simplicity means it can rapidly fit data on new and emerging pathogens. A key component of our modeling framework is the inclusion of farm-infection status in the calculations of the local-transmission and network effects. We found this inclusion consistently improved the fit of our models no matter the criterion used. Our findings imply that what matters is not being near other farms but being near other infected farms; likewise, what matters is not receiving pigs from another farm but receiving infected pigs. Thus, our results demonstrated that both effects should be considered.
